# The nidogen-domain containing protein DEX-1 is required for epidermal remodeling in *C. elegans* dauers

**DOI:** 10.1101/277798

**Authors:** Kristen M. Flatt, Caroline Beshers, Cagla Unal, Nathan E. Schroeder

## Abstract

Phenotypic plasticity is a critical component of an organism’s ability to thrive in an ever-changing environment. The free-living nematode, *Caenorhabditis elegans*, adapts to unfavorable environmental conditions by pausing reproductive development and entering a stress-resistant larval stage known as dauer. The transition into dauer is marked by vast morphological changes – including remodeling of epidermis, neurons and muscle. Though many of these dauer-specific traits have been described, the molecular basis of dauer-specific remodeling is still poorly understood. Here we show that the nidogen-domain containing protein DEX-1 functions downstream of the dauer decision to facilitate stage-specific tissue remodeling during dauer morphogenesis. DEX-1 was previously shown to regulate sensory dendrite formation during embryogenesis. We find that DEX-1 is also required for the proper remodeling of the stem cell-like epidermal seam cells and maintenance of seam cell quiescence during dauer. *dex-1* mutant dauers lack distinct lateral cuticular alae during dauer and have increased sensitivity to sodium dodecyl sulfate (SDS). Furthermore, we find that DEX-1 mediated seam cell remodeling is required for proper dauer mobility. We show that DEX-1 acts cell autonomously in the seam cells during dauer and that *dex-1* expression during dauer is regulated through DAF-16/FOXO-mediated derepression. Finally, we show that *dex-1* interacts with a family of zona pellucida-domain encoding genes to regulate dauer-specific epidermal remodeling. Taken together, our data indicates that DEX-1 plays a central role in *C. elegans* epidermal remodeling during dauer.

## Introduction

In order to survive changing environments, organisms modify their phenotype (i.e. phenotypic plasticity). Tissue remodeling is an important component of stress-induced phenotypic plasticity. For example, desert locusts are capable of altering their morphology between distinct ‘gregarious’ and ‘solitarious’ phases based on population density (Pener and Simpson 2009), and certain species of butterfly also change their body morphology and wing color based on seasonal cues (Windig 1994). While these large-scale displays of phenotypic plasticity are readily observed, the molecular basis of tissue remodeling in response to environmental inputs is often unclear.

*C. elegans* is a useful animal to investigate the molecular mechanisms that facilitate stress-induced remodeling. Under favorable growth conditions, *C. elegans* develops continuously through four larval stages (L1-L4) into a reproductive adult. However, under unfavorable environmental conditions, *C. elegans* larvae arrest their development at the second larval molt and enter the stress-resistant dauer stage (Cassada and Russell 1975; Golden and Riddle 1984). Dauers are specialized, non-feeding larvae capable of withstanding extended periods of adverse environmental conditions. Dauer-specific stress resistance is likely facilitated by several morphological changes that occur during dauer formation. For example, dauers display both structural and biochemical differences in their epidermis and cuticle compared with non-dauers (Cox *et al.* 1981; Blaxter 1993). Dauer formation also corresponds with a general radial shrinkage of the body and the formation of longitudinal cuticular ridges called alae (Fig. 1A).

**Figure 1.**
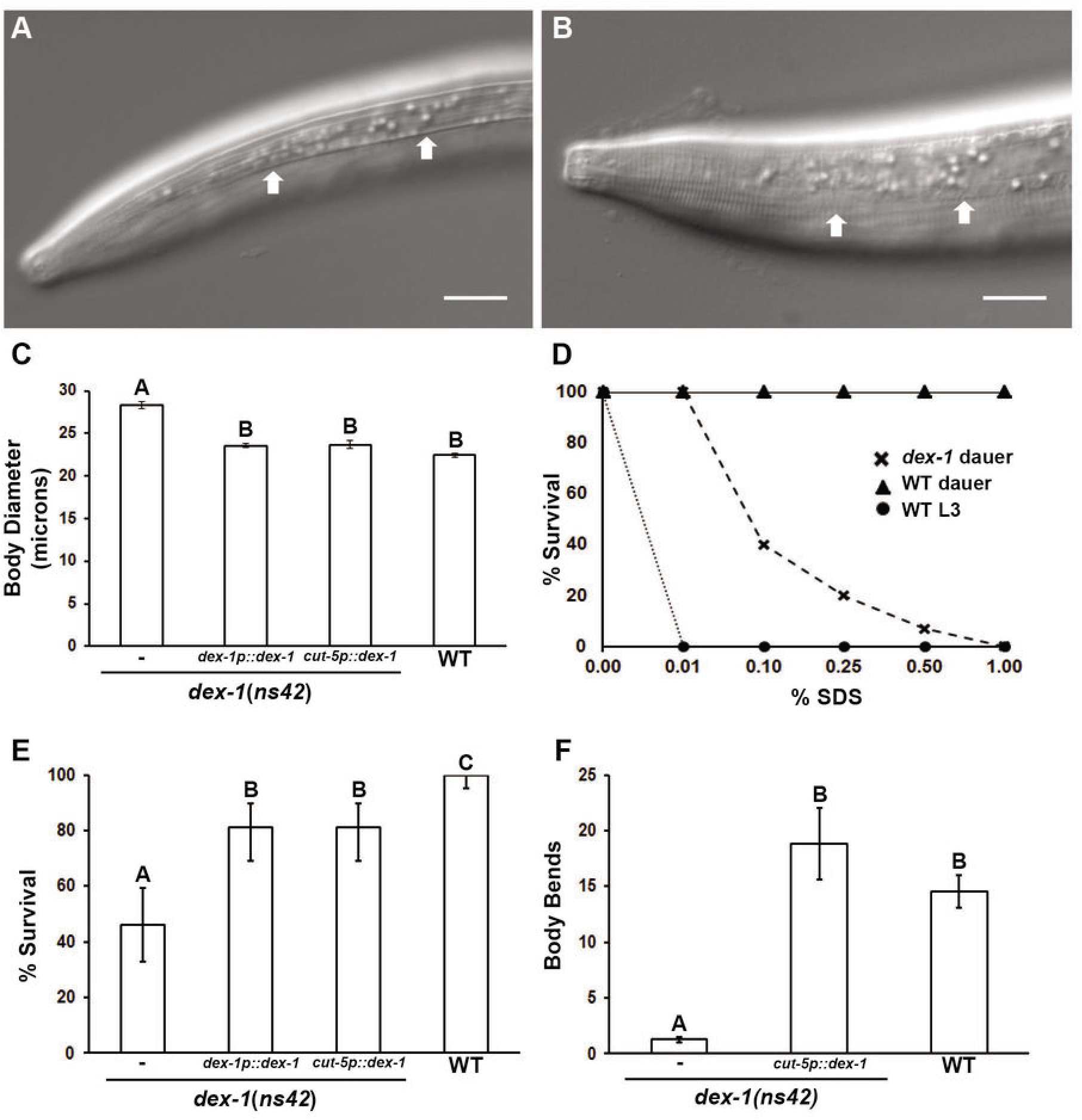
*dex-1* mutants form partial dauers. Wild-type dauers (A) have prominent lateral alae, while *dex-1* mutant dauers’ (B) alae are indistinct. Arrows point to the position of the lateral alae. Scale bar, 10μm. (C) *dex-1* mutant dauers are defective for dauer-specific radial shrinkage. The radial shrinkage defect can be rescued from both the endogenous *dex-1* promoter and a seam cell-specific promoter (*cut-5p::dex-1*).Error bars, SEM. Different letters above bars indicate statistically significant differences (α = 0.05, n = 20). (D) Dose response survival assay to SDS. *dex-1* dauers are sensitive to SDS compared to wild-type dauers, but are able to survive low levels of SDS exposure. Non-dauer animals are sensitive at all tested SDS concentrations (n = 60).(E) Percent survival of dauers to 0.1 % SDS. *dex-1* mutant dauer SDS sensitivity phenotype was rescued using both endogenous *dex-1* and a seam cell-specific promoter. Data was analyzed using a one proportion exact method analysis. Error bars, 95% C.I. (n = 60). (G) *dex-1* mutant dauers are sluggish when mechanically stimulated. Expression of *dex-1* from the seam cells rescued the locomotion response to wild-type levels. Data were analyzed by Kruskal-Wallis test with Dunn’s multiple comparisons test (α = 0.05, n = 40). Error bars, SEM.

Radial shrinkage and alae formation are regulated by a set of lateral hypodermal seam cells (Singh and Sulston 1978; Melendez *et al.* 2003). Seam cell function and remodeling are critical for proper dauer morphology and increased environmental resistance. The seam cells also have stem cell-like properties. During non-dauer development, the seam cells undergo asymmetrical divisions at larval molts to produce an anterior differentiated cell and a posterior seam cell. Alternatively, if the animal enters dauer diapause, the seam cells shrink and stop dividing.

Here, we characterize the role of DEX-1, a protein similar to mammalian tectorin and SNED1, in remodeling of the seam cells during dauer formation. DEX-1 comprises a transmembrane protein with two extracellular nidogen domains that is required for proper sensory neuron dendrite formation during embryogenesis (Heiman and Shaham 2009). We find that DEX-1 is also required during dauer formation for seam cell remodeling and resistance to environmental stressors. Furthermore, we find that *dex-1* is upregulated in seam cells during dauer in a DAF-16 dependent manner. DEX-1 was previously shown to interact with a zona pellucida (ZP) domain containing protein to mediate dendrite extension. Our data suggest that DEX-1 interacts with additional ZP-domain proteins, to regulate seam cell remodeling. Finally, we implicate DEX-1 in regulating seam cell quiescence. Combined with previous data demonstrating a role for DEX-1 in sensory dendrite adhesion (Heiman and Shaham 2009), our data suggest that DEX-1 plays a role in modulating cell shape of several cell types throughout development.

## Materials and Methods

### Strains

All strains were grown under standard conditions unless otherwise noted (Brenner 1974). The wild-type Bristol N2 strain and the following mutant strains were used: CHB27 *dex-1(ns42)* III; SP1735 *dyf-7(m537)* X; CB1372 *daf-7(e1372)* III; DR129 *daf-2(e1370) unc-32(e189)* III; DR27 *daf-16(m27)* I; FX01126 *cut-1(tm1126)* IIl; RB1574 *cut-6(ok1919)* III; RB1629 *cut-5(ok2005)* X. *dex-1(ns42)* was given to us by Dr. Maxwell Heiman (Department of Genetics, Harvard University). *cut-1(tm1126)* was provided by the Mitani Consortium (Department of Physiology, Tokyo Women’s Medical University School of Medicine, Japan). All other mutant strains were provided by the Caenorhabditis Genetics Center (CGC), which is funded by NIH Office of Research Infrastructure Programs (P40 OD010440). The following transgenic animals were also provided by the CGC: ST65 *ncIs13[ajm-1::GFP]*; JR667 *wIs51 [SCMp::GFP+unc-119(+)]*. The IL2 neurons were observed using PT2660 *myIs13[klp-6p::GFP+pBx]* III (Schroeder et al., 2013); PT2762 *myIs14[klp-6p::GFP +pBx]* V and JK2868 *qIs56*[*lag-2p::GFP*] (Blelloch *et al.*; Ouellet *et al.* 2008; Schroeder *et al.* 2013).

The following plasmids were the generous gift of Dr. Maxwell Heiman: pMH7 *dex-1p::dex-1*, pMH8 *pha-4p::dex-1*, pMH111 *dex-1p(5.7kb)::gfp*, pMH125 *dex-1p(2.1kb)::gfp* (Heiman and Shaham 2009). Novel plasmids were constructed using Gibson Assembly (NEB, E2611S). The seam cell-specific expression plasmid was built by replacing the *dex-1* promoter from pMH7 with a 1.21kb *cut-5* promoter region (for a complete list of primers used to construct plasmids, see Table S1). The hypodermal-specific *dex-1* plasmid was constructed by replacing the *dex-1* promoter in pMH7 with the *dpy-7* promoter (Gilleard *et al.* 1997). The IRS sequence was deleted from pMH111 using the Q5 Site Directed Mutagenesis Kit (NEB, E05525).

Animals containing extrachromosomal arrays were generated using standard microinjection techniques (Mello *et al.* 1991) and genotypes confirmed using PCR analysis and observation of co-injection markers. Each animal was injected with 20μL of plasmid and 80μL of *unc-122p::gfp* as the co-injection marker.

### Dauer Formation

Dauers were induced by one of two methods. For non-temperature sensitive strains, we used plates containing crude dauer pheromone extracted by previously established procedures (Vowels and Thomas 1992; Schroeder and Flatt 2014). For animals with mutations in *daf-7(e1372)* or *daf-2(e1370)*, dauers were induced using the restrictive temperature of 25°C (Riddle *et al.* 1981).

### Microscopy and Rescue Analysis

Animals were mounted onto 4% agarose pads and anesthetized with 1.0 mM or 0.1 mM levamisole for dauers and non-dauer or partial dauers, respectively. A Zeiss AxioImager microscope equipped with DIC and fluorescent optics was used to collect images. Images were analyzed using FIJI. For radial constriction experiments, measurement data was analyzed using a One-Way ANOVA with Bonferroni’s multiple comparisons test using Graphpad Prism 6 software. The seam cell area was measured for V2pap, V2ppp and V3pap and averaged to give one measurement per animal (Sulston and Horvitz 1977).

### SDS Sensitivity Assays

Sodium dodecyl sulfate (SDS) dose-response assays were performed using 12-well culture dishes with wells containing M9 buffer and specified concentrations of SDS dissolved in water. Twenty dauers from each treatment were transferred into SDS. Dauer animals were exposed to SDS for 30 minutes and scored using established criteria (Liu *et al.* 2013) Briefly, animals were stimulated once with an eyelash and were scored as alive if movement was observed. Each concentration was tested in triplicate with each experiment containing a separate wild-type (N2) control. The LD_50_ and confidence interval of each concentration was calculated using probit analysis in Minitab 18.

### Fluorescent Bead Feeding Assay

Feeding assays were carried out using established methods (Nika *et al.* 2016).Briefly, fluorescent beads (Sigma L3280) were added to a 10X concentrated OP50 *E. coli* overnight culture. Fresh NGM plates were then seeded with 65 µL of the bead/bacteria suspension and allowed to dry. Twenty dauer or non-dauer animals were added to the plate and incubated at 20°C for 40 minutes. Worms were then observed for the presence of fluorescent beads in the intestinal tract. Each experiment was performed twice.

### Locomotion Assays

Animals were transferred to unseeded NGM plates and allowed to sit at room temperature for 10 minutes before being assayed. Animals were stimulated near the anus with an eyelash and the number of body bends was scored. Counting was stopped if the animal did not complete another body bend within 5 seconds of stopping, or if the animal reversed direction. Each animal was scored twice and then removed from the plate. Counts were averaged and then analyzed using a Kruskal-Wallis test with Dunn’s multiple comparisons test for dauers and the Mann-Whitney U test for adult animals using Graphpad Prism 6 software.

### DEX-1 Expression Analysis

The fluorescent intensities of the V2pap, V2ppp and V3pap seam cells were measured using established methods (McCloy *et al.* 2014). Briefly, each cell was outlined and the area, integrated density and mean gray value were measured. Measurements were also taken for areas without fluorescence surrounding the cell. The total corrected cell fluorescence (TCCF = integrated density - (area of selected cell * mean fluorescence of background reading)) was then calculated for each cell. The intensities of the three cells from each worm were averaged such that each nematode comprised a single data point. The data were analyzed using One-Way ANOVA and Bonferroni’s multiple comparisons test. Ten animals were measured for each genotype.

## Results

### DEX-1 is required for proper dauer morphology and behavior

Wild-type *C. elegans* dauers have a distinctive morphology due to radial shrinkage that leads to a thin appearance compared with non-dauers (Figure 1A). We found that *dex-1(ns42)* mutants are defective in dauer radial shrinkage (Figure 1C). *dex-1* mutant dauers are significantly wider in diameter when compared to wild-type dauers, leading to a ‘dumpy dauer’ phenotype (Figure 1, B and C). The defect in body size appears specific to the dauer stage, as non-dauer *dex-1* mutants show no differences in body size compared with wild-type non-dauers (Figure S1). Radial shrinkage in dauers is correlated with the formation of longitudinal cuticular ridges on the lateral sides of the animal called the alae (Cassada and Russell 1975). The lateral alae of *dex-1* mutant dauers are indistinct compared with wild-type dauers (Figures 1B and 2S). To confirm *dex-1* as the causative mutation, we rescued the radial shrinkage phenotype of *dex-1* mutants with *dex-1* cDNA under the control of its endogenous promoter (Figure 1C). Together, these data suggest that *dex-1* mutants form partial dauers with defects in epidermal remodeling.

In addition to radial shrinkage, dauers have several structural modifications compared with non-dauer animals that lead to increased resistance to environmental insults (Cassada and Russell 1975; Androwski *et al.* 2017). We characterized several dauer-specific phenotypes in *dex-1* mutants. First, we exposed dauer animals to SDS. Wild-type dauers survive for hours in 1% SDS (Cassada and Russell 1975). We found that while *dex-1* mutant dauers were sensitive to 1% SDS, they were able to survive significantly higher concentrations of SDS that wild-type non-dauer animals (Figure 1D) Similar to our radial shrinkage data, we could effectively rescue the SDS phenotype with a wild-type copy of *dex-1* (Figure 1E). Second, we used a fluorescent bead assay that determines whether feeding occurs (Nika *et al.* 2016). Non-dauers will readily ingest beads and show fluorescence throughout the digestive system. Dauers suppress pharyngeal pumping and have a buccal plug that prevents entry of fluorescent beads. We found that while fluorescence was never observed in *dex-1* mutant dauer intestines, we occasionally observed fluorescence in the buccal cavity (Figure S3). We never observed pharyngeal pumping in *dex-1* dauers. These data suggest that while pharyngeal pumping is efficiently suppressed, *dex-1* dauers have low-penetrance defects in buccal plug formation. Finally, we examined *dex-1* dauers for the presence of dauer-specfic gene expression of *lag-*2p::*gfp* in the IL2 neurons during dauer (Ouellet *et al.* 2008). Similar to wild-type dauers, *dex-1* dauers showed appropriate dauer-specific expression, indicating that they are indeed dauers with defects limited to epidermal remodeling (data not shown).

The decision to enter dauer is based on the ratio of population density to food availability (Golden and Riddle 1984). *C. elegans* constitutively secrete a pheromone mixture that is sensed by animals and, at high levels, triggers dauer formation (Golden and Riddle, 1982). Dauers can be picked from old culture plates (starved) or can be induced using purified dauer pheromone. We found no difference in the dauer phenotype between starved or pheromone-induced *dex-1* mutant dauers. The *C. elegans* insulin/IGF-like and TGF-β signaling pathways function in parallel to regulate the dauer formation decision. Reduced insulin and TGF-β signaling induced by overcrowding and scarce food promotes dauer formation. Disruption of either the insulin-receptor homolog DAF-2 or the TGF-β homolog DAF-7 results in constitutive formation of dauers. Double mutants of *daf-2* or *daf-7* with *dex-1* did not suppress the defects in dauer morphogenesis of *dex-1*, suggesting that *dex-1* is acting downstream of the dauer decision pathway (data not shown).

### DEX-1 functions in the stem cell-like seam cells to regulate dauer morphogenesis

*dex-1* is expressed at high levels during embryogenesis to regulate sensory dendrite formation (Heiman and Shaham 2009). However, previous microarray data suggested that *dex-1* is also upregulated during dauer (Liu *et al.* 2004). We generated transgenic animals expressing GFP driven by a 5.7kb 5’ *dex-1* upstream promoter and observed bright fluorescence in the seam cells and glia socket cells of the anterior and posterior deirid neurons starting in the pre-dauer L2 (L2d) stage. Expression of *dex-1p::gfp* in the seam cells and deirid socket cells persists throughout dauer and is downregulated upon recovery from dauer (Figure 2, A and B). We also observed weak *dex-1p::gfp* expression in unidentified pharyngeal cells during all larval stages (Figure S4).

**Figure 2.**
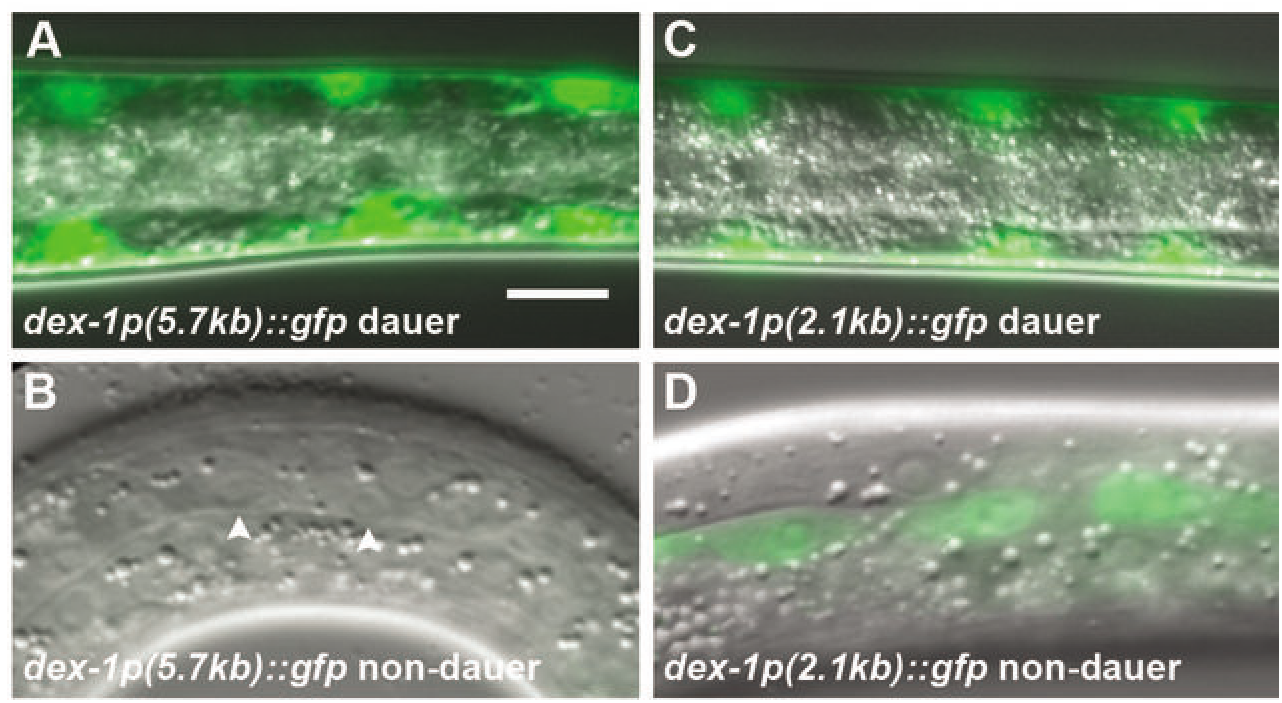
The 5.7 kb upstream promoter region of *dex-1* contains dauer-specific regulatory elements. Dorsoventral view of (A) dauer and lateral view of (B) non-dauer animals expressing GFP from a 5.7kb upstream promoter region of *dex-1*. Fluorescence in the seam is observed exclusively during dauer (Arrowheads indicate the seam cells). Dorsoventral view of a (C) dauer and lateral view of (D) non-dauer expressing GFP from a 2.1kb upstream promoter region. Using the short promoter, GFP expression is observed in the seam cells during all larval stages including dauer. Scale bar, 10μm.

Dauer-specific radial shrinkage and subsequent lateral alae formation are facilitated by shrinkage of the seam cells (Singh and Sulston, 1978; Melendez et al, 2003). Using the *ajm-1*::*gfp* apical junction marker (Koppen et al., 2001), we found that *dex-1* mutant dauer seam cells are larger and have jagged, rectangular edges unlike the smooth, elongated seam cells of wild type dauers (Figure 3). These data suggest that DEX-1 is required for seam cell remodeling. The seam cells also have stem cell-like properties. During non-dauer development, seam cells divide at larval molts to produce a seam cell daughter and a differentiated daughter cell (Sulston and Horvitz, 1977). During dauer, seam cells enter a quiescent state and only resume division following recovery from dauer. To determine if *dex-1* is required for maintaining seam cell quiescence during dauer, we used a seam cell nuclei marker to examine the number of seam cell nuclei in wild-type and *dex-1* backgrounds. We found that *dex-1* mutant dauers have a slight, but statistically significant, greater number of seam cell nuclei compared with wild type dauers (*dex-1* 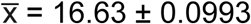, WT 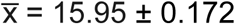, p = 0.0010, n = 40). This could suggest that DEX-1 plays a role in maintaining seam cell quiescence during dauer.

**Figure 3.**
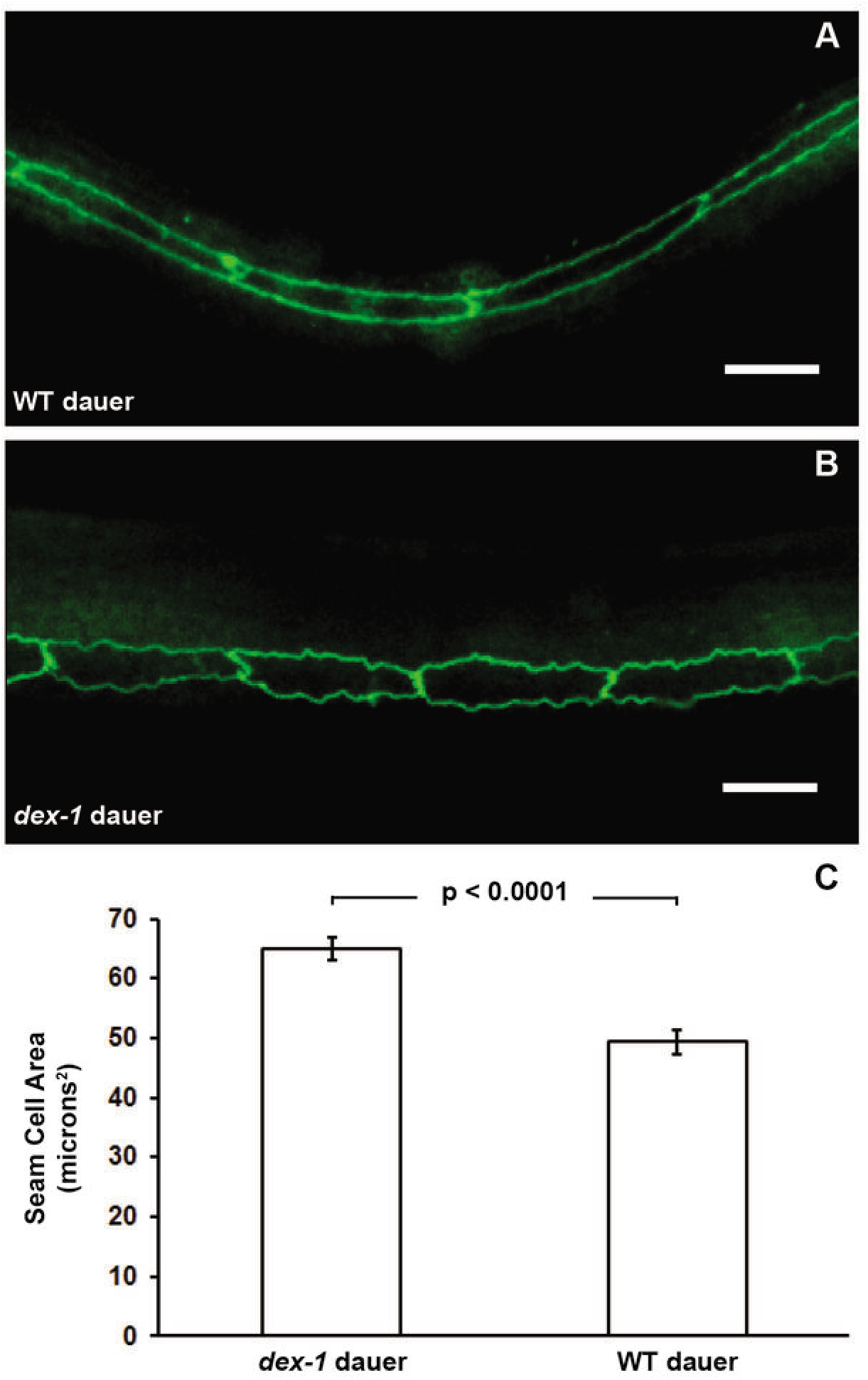
*dex-1* mutant dauers have defects in seam cell shrinkage. Lateral view micrographs of (A) wild type and (B) *dex-1* dauers expressing the apical junctions marker *ncIs13* [*ajm-1::gfp*]. The seam cells of wild-type dauers are elongated and smooth, while *dex-1* mutant seam cells are rectangular with jagged edges. Scale bar, 10μm.(C) Quantification of seam cell area as measured with the *ajm-1::gfp* reporter demonstrates that seam cells in *dex-1* mutant dauers are also significantly larger than in wild-type dauers. Data were analyzed by an unpaired t-test (*dex-1* dauer n = 14, WT dauer n = 15). Error bars, SEM.

To determine where *dex-1* acts to regulate seam cell remodeling, we expressed *dex-1* cDNA under the control of cell-specific promoters. First, we expressed *dex-1* in the seam cells using the *cut-5* promoter. *cut-5* was previously shown to be expressed specifically in the seam cells during L1 and dauer (Sapio *et al.* 2005). Using this seam cell-specific promoter, we partially rescued SDS sensitivity in *dex-1* mutant dauers (Figure 1E). The seam cell-specific expression of *dex-1* resulted in a mosaic rescue of the radial shrinkage phenotype (Figure S5). The rescued seam cells that underwent radial constriction also displayed intact lateral alae, while the non-rescued seam cells’ alae remained indistinct. Expression of *dex-1* under a pharyngeal promoter failed to rescue the *dex-1* seam cell phenotype suggesting a cell autonomous role for DEX-1. To verify the cell autonomous nature of *dex-1*, we sought to examine expression in a tissue with close contact to the seam cells. The seam cells are surrounded by a syncytial hypodermis. *dex-1* expression in the surrounding hypodermis failed to rescue the *dex-1* dauer-specific phenotype. Interestingly, expression of *dex-1* in the hypodermis induced a dumpy phenotype in approximately 33% of non-dauer animals (Fig. S6). Together, these data indicate that *dex-1* acts cell-autonomously to regulate seam cell remodeling during dauer.

### DEX-1 acts in seam cells to regulate locomotion during dauer

Morphological changes during dauer are accompanied by changes in behavior. Wild-type dauer animals are general quiescent, but move rapidly when mechanically stimulated (Cassada and Russell 1975). Anecdotally, we noticed a higher percentage of quiescent *dex-1* dauers than seen with wild-type dauers. To quantify this behavior, we developed a behavioral assay that quantifies movement following mechanical stimulation (see Materials and Methods). Although both *dex-1* mutant and wild-type dauers initially respond to mechanical stimulation, the *dex-1* mutant dauers have significantly reduced locomotion and display slightly uncoordinated body movements (Figure 1F and data not shown). This locomotion defect was dauer-specific, as non-dauer *dex-1* animals moved at wild-type levels following mechanical stimulation (Figure S1).

Given that *dex-1* was primarily expressed in the seam cells during dauer, we tested if seam cell-specific expression could rescue the behavioral phenotype. Surprisingly, seam cell-specific *dex-1* expression completely rescued the *dex-1* dauer locomotion defects (Figure 1F). In addition to seam cell remodeling, several neuron types remodel during dauer formation (Albert and Riddle 1983). For example, the IL2 and deirid sensory neurons remodel during dauer formation (Albert and Riddle 1983; Schroeder *et al.* 2013). The IL2s regulate dauer-specific behaviors (Lee *et al.* 2011; Schroeder *et al.* 2013), while the deirids respond to specific mechanical cues (Sawin *et al.* 2000). We, therefore, examined these neuron classes using fluorescent reporters; however, we observed no obvious difference in the neuronal structures between *dex-1* and wild-type (data not shown).

### *dex-1* expression in dauers is regulated by DAF-16

To understand how *dex-1* expression is regulated, we examined the 5’ upstream region of *dex-1* for potential regulatory sites. Previous Chromatin Immunoprecipitation (ChIP-seq) data identified a putative DAF-16 binding site upstream of the *dex-1* coding region (Celniker *et al.* 2009). DAF-16 is the sole *C. elegans* ortholog of the human Forkhead BoxO-type transcription factor and a major regulator of the dauer decision. To examine whether this region affects expression of *dex-1*, we first expressed GFP from a truncated 2.1kb *dex-1* promoter. Unlike the 5.7kb *dex-1* promoter fusion, the short *dex-1* promoter drove GFP expression in the seam cells during all larval stages (Figure 2, C and D). This suggests there are elements within the long promoter that repress *dex-1* expression during non-dauer stages. While mutations in *daf-16* result in animals incapable of forming dauers, under conditions of high pheromone concentrations *daf-16* mutants can enter into a partial dauer state (Vowels and Thomas 1992; Gottlieb and Ruvkun 1994). *daf-16* partial dauers are identifiable by body morphology and the presence of indistinct lateral alae (Vowels and Thomas 1992). To determine if DAF-16 is regulating *dex-1* expression during dauer, we first examined the expression of the 5.7 kb *dex-1p::*gfp reporter in *daf-16* partial dauers. We found that the fluorescent intensity of *dex-1p::gfp* was significantly reduced in the *daf-16* partial dauers compared to wild-type, suggesting that DAF-16 regulates *dex-1* seam cell expression during dauer (Figure 4, B-D).

**Figure 4.**
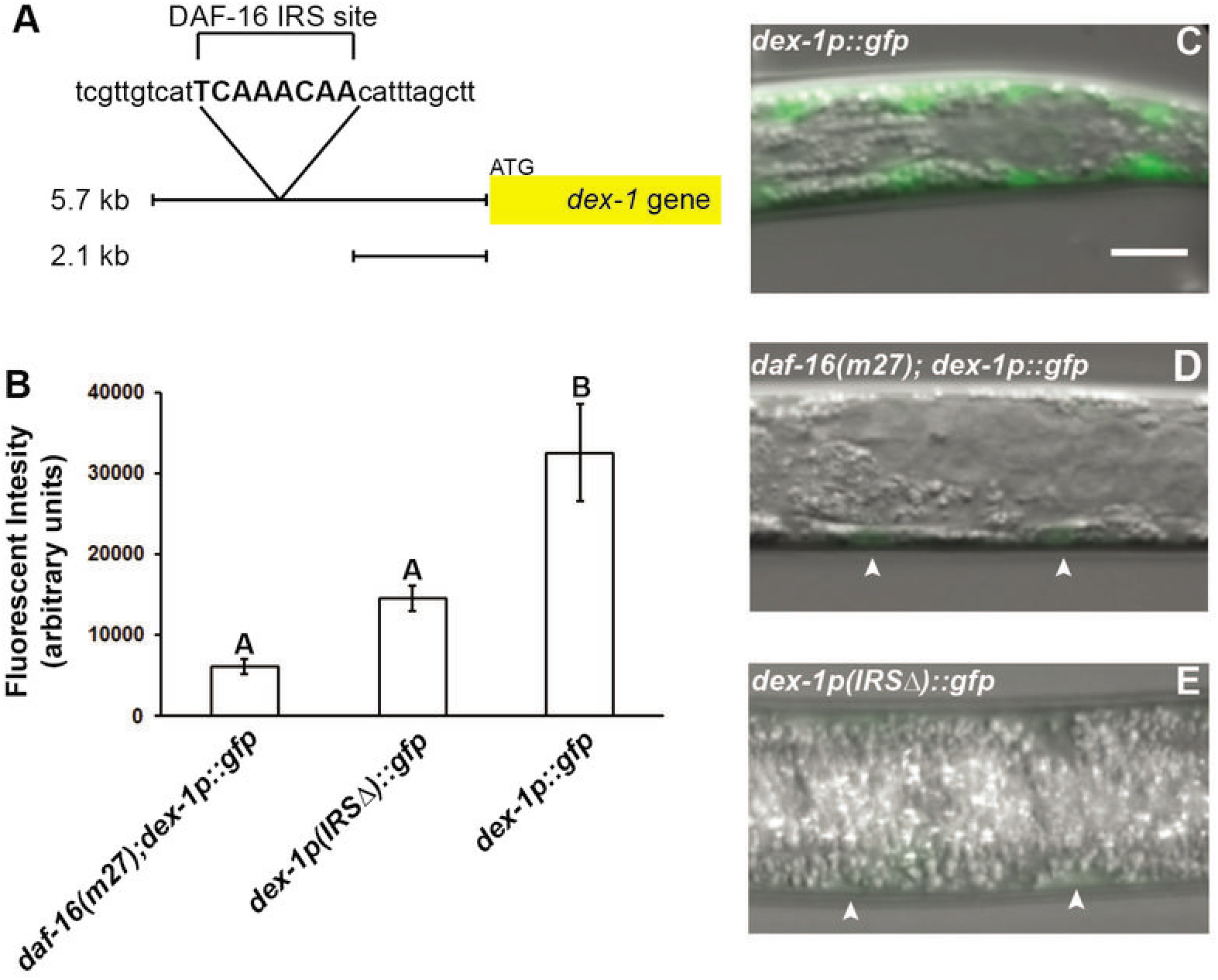
*dex-1* expression in the seam cells is regulated by DAF-16. (A) We identified a putative DAF-16 IRS binding site (in all caps) ∼2.8 kb upstream from the *dex-1* ATG start site. (B) Quantification and (C-E) dorsoventral view micrographs of GFP expression driven by the 5.7 kB *dex-1* promoter in dauers. Expression of GFP from the 5.7 kB *dex-1* promoter in a (C) wild-type background produces bright fluorescence in the seam cells during dauer (Also see Figure 2A). Fluorescent intensity is reduced in (D) *daf-16*(*m27)* partial dauer mutant background. Deletion of the DAF-16 IRS sequence (E) from the 5.7kb promoter region also significantly reduces GFP expression in the seam cells during dauer. Arrowheads indicate seam cells. Scale bar, 10μm. Different letters above graph indicate statistically significant differences using a One-way ANOVA with Bonferroni’s multiple comparisons correction (α = 0.05, n = 10). Error bars represent the SEM.

FOXO/DAF-16 binds to canonical DAF-16 binding elements and insulin response sequences (IRS) (Paradis and Ruvkun 1998; Obsil and Obsilova 2008). Within the ChIP-seq identified region (Celniker *et al.* 2009), we identified a putative insulin response sequence (IRS) binding site (Figure 4A). To determine whether the identified DAF-16 IRS site directly regulates *dex-1* expression, we deleted the IRS sequence in the 5.7kb *dex-1* promoter region used to drive GFP. We found that deleting the IRS sequence results in reduced GFP expression in the seam cells during dauer, similar to the levels observed in *daf-16* partial dauers (Figure 4, B-E). Taken together, these results indicate that a repressor element lies within the 5.7kb promoter and that DAF-16 binding to the IRS acts to derepress *dex-1* expression during dauer formation.

### *dex-1* interacts with genes encoding ZP-domain proteins

DEX-1 interacts with the ZP-domain protein DYF-7 to regulate primary dendrite extension during embryogenesis (Heiman and Shaham 2009). We, therefore, examined the *dyf-7(m537)* mutant for defects in dauer morphogenesis. Unlike *dex-1* mutants, *dyf-7* mutants are defective for dauer formation under typical dauer-inducing environmental conditions (Starich *et al.* 1995). We, therefore, examined *dyf-7* mutants in dauer formation constitutive mutant backgrounds. Both *daf-2; dyf-7* and *daf-7; dyf-7* double mutants were normal for dauer-specific radial shrinkage, SDS resistance, and IL2 arborization (data not shown).

The cuticlin (CUT) proteins are a family of ZP-domain proteins including several that are required for proper dauer morphology. Disruption of *cut-1, cut-5* and *cut-6* result in dauers with incomplete radial shrinkage and defective alae formation (Sebastiano *et al.* 1991; Muriel *et al.* 2003; Sapio *et al.* 2005b). We asked whether these defects in CUT mutant larvae were due to seam cell remodeling. We found that, similar to *dex-1*, the seam cells of the CUT mutants were rectangular with jagged edges (Figure 5). Also similar to *dex-1*, we found that *cut-1* and *cut-5* dauers were more sensitive to SDS compared with wild type dauers (Table 1). Interestingly, while *cut-6* mutant dauers were resistant to the standard 1% SDS treatment (Muriel *et al.* 2003), we found that the *cut-6* mutant dauers were substantially more sensitive to SDS than wild-type dauers (Table 1). Taken together, these data indicate similar roles for DEX-1 and cuticlins during dauer remodeling.

**TABLE 1:** SDS concentration necessary to kill 50% of animals tested for each genotype.

| STRAIN | LD<sub>50</sub><sup>a</sup> |
| --- | --- |
| WT dauer | >10% ± --- |
| WT L3 | <0.01 ± --- |
| *dex-1(ns42)* | 0.15 ± 0.03 |
| *cut-1(tm1126)* | 0.28 ± 0.05 |
| *cut-6(ok1919)* | 2.02 ± 0.15 |
| *cut-5(ok2005)* | 0.25 ± 0.06 |
| *cut-1;dex-1* | 0.20 ± 0.03 |
| *dex-1;cut-6* | 0.56 ± 0.29 |
| *dex-1;cut-5* | ---<sup>b</sup> |
| *cut-1;cut-5* | 0.08 ± 0.02 |
| *cut-1;cut-6* | 1.85 ± 0.10 |
| *cut-6;cut-5* | 0.01 ± ---<sup>c</sup> |

<sup>a</sup> 50% lethal dose ± 95% confidence interval
<sup>b</sup> Synthetic lethal at L1
<sup>c</sup> Synthetic dumpy at all stages

**Figure 5.**
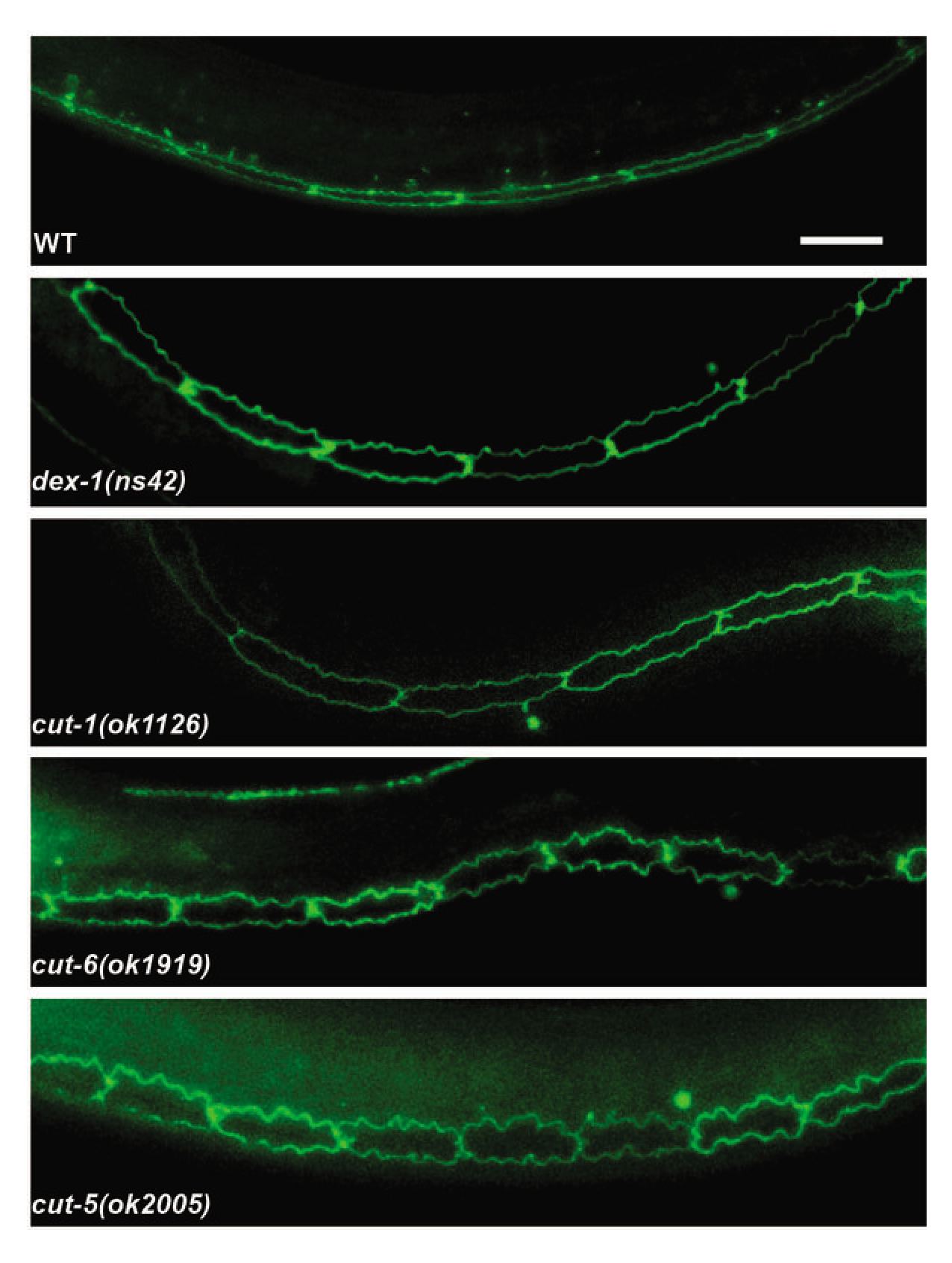
Cuticlin mutants phenocopy the *dex-1* mutant seam cell phenotype during dauer. Lateral view of wild-type, *dex-1* and cuticlin mutant dauers expressing the apical junctions marker *ajm-1p::GFP*. The seam cells in cuticlin mutants are jagged and rectangular, closely resembling those of *dex-1* mutant dauers. Scale bar, 10μm.

We hypothesized that, similar to its interaction with DYF-7 during embryogenesis, DEX-1 may interact with the CUT proteins during dauer. We, therefore, examined double mutants of *dex-1* with *cut-1, cut-5*, and *cut-6.* The *dex-1; cut-1* double mutant did not enhance SDS sensitivity beyond the *dex-1* single mutant, suggesting that they may act in the same pathway to regulate dauer remodeling. The *dex-1; cut-5* double mutant was synthetically lethal during embryogenesis or early L1. This is similar to the *dex-1; dyf-7* double mutant (Heiman and Shaham 2009), suggesting that in addition to roles in dauer remodeling, *cut-5* and *dex-1* have redundant roles during embryogenesis. However, unlike the *dex-1; dyf-7* double mutant (Heiman and Shaham 2009), the *dex-1; cut-5* synthetic lethality did not display temperature sensitivity (data not shown). Interestingly, the *dex-1cut-6* double mutant was intermediate in SDS sensitivity between the *dex-1 and cut-6* single mutants (Table 1).

We further tested the *cut* mutant phenotypes by generating double mutants between each of the *cut* mutants (Table 1). The *cut-1;cut-5* double mutant showed a significant reduction of SDS resistance compared to single mutants alone, suggesting that they may be acting in parallel pathways. The *cut-1;cut-6* double mutants retained the *cut-6* SDS sensitivity phenotype, suggesting that *cut-6* is epistatic to *cut-1.* The *cut-6;cut-5* dauers showed a drastic increase in sensitivity to SDS compared to the single mutants, indicating that these genes may also play roles in parallel pathways during dauer remodeling. Interestingly, the *cut-6; cut-*5 double mutant showed a severe dumpy phenotype in all developmental stages. These results confirm previous work demonstrating a broad role for ZP-domain proteins in development (Heiman and Shaham 2009; Gill *et al.* 2016).

## Discussion

The dauer stage of *C. elegans* is an excellent example of a polyphenism, where distinct phenotypes are produced by the same genotype via environmental regulation (Simpson *et al.* 2011). Compared to the decision to enter dauer, little is known about the molecular mechanisms controlling remodeling of dauer morphology. DEX-1 was previously characterized as a component of embryonic neuronal development (Heiman and Shaham 2009). Our data shows that DEX-1 also functions during dauer-specific remodeling of the stem cell-like seam cells. We demonstrate a cell-autonomous role for DEX-1 in the regulation of seam cell remodeling during dauer morphogenesis.

We found that *dex-1* is required for the shrinkage of the seam cells and formation of lateral alae during dauer. Furthermore, we found that *dex-1* dauers have significantly more seam cells than wild-type dauers, suggesting ectopic divisions during dauer diapause. In mammalian cell lines, stem cell shape regulates differentiation (McBeath *et al.* 2004; Kilian *et al.* 2010). For example, mesenchymal stem cells will differentiate into adipocytes or osteoblasts depending on whether the cell is round or flat, respectively (McBeath *et al.* 2004). We speculate that the shrinkage observed during dauer may be important for maintenance of seam cell quiescence.

We also found the *dex-1* mutant dauers have defects in locomotion when mechanically stimulated. We originally assumed that this could be due to the lack of lateral alae; however, our seam-cell specific rescue of *dex-1* resulted in a mosaic pattern of alae formation while having complete rescue of behavior. We, therefore, speculate that *dex-1* mutant dauers may have defects in other neuron classes that result in locomotion defects during dauer. Previous RNAi data shows that knockdown of *dex-1* results in low penetrance defects in motor neuron commissure formation (Schmitz *et al.* 2007). Dauers exhibit changes in somatic muscle structure (Dixon *et al.* 2008). It will be interesting to determine if *dex-1* mutants have dauer-specific defects in neuromuscular junctions.

Dissection of the genetic pathways regulating the decision to enter dauer has revealed insights into TGF-β, insulin and hormone signaling. The FOXO transcription factor, DAF-16, is a well-known regulator of the dauer formation decision by acting downstream of the insulin/IGF-1 receptor DAF-2. Dauer-inducing environmental conditions lead to a translocation of DAF-16 to the nucleus where it activates dauer formation pathways (Lee *et al.* 2001; Fielenbach and Antebi 2008). We found that *dex-1* expression during dauer is regulated by DAF-16. Based on our results we propose that DEX-1 is normally repressed during non-dauer post-embryonic stages and DAF-16 serves to derepress *dex-1* expression via an upstream insulin response sequence. The *dex-1p::gfp* fluorescence was not completely eliminated in the *daf-16* partial dauers and the IRS-deletion dauers suggesting that both *dex-1* repressors and other regulatory elements remain to be identified.

Together with previous work (Heiman and Shaham 2009), our data suggests that DEX-1 may interact with ZP-domain proteins. While the *cut* mutants are all deletion alleles that disrupt the ZP-domains and, therefore, likely functional nulls, the sole *dex-1(ns42)* allele is a nonsense mutation that disrupts a region resembling mammalian zonadhesin (Heiman and Shaham 2009). It is possible that *dex-1* is a non-null and, therefore, genetic interaction experiments should be interpreted with caution. It has been proposed that biochemical compaction of the cuticlins in the extracellular space between the seam and the hypodermis causes radial constriction, and thus forms the lateral alae via a ‘cuticlin tether’ (Sapio *et al.* 2005). We add to this by proposing that DEX-1 may function as a molecular binding hub for multimerized cuticlins. Our data indicate that DEX-1 acts in a cell autonomous manner within the seam cells. DEX-1 includes a transmembrane domain and, therefore, may function as a transmembrane anchor while simultaneously binding the multimerized cuticlins in the extracellular space (Figure 6). Alternatively, DEX-1 could act as a secreted factor with restricted localization to the apical border of the seam cell. During embryogenesis, DEX-1 is secreted and localized to the dendritic tips (Heiman and Shaham, 2009). In both models, DEX-1 may serve to couple physical interactions between the remodeled cuticular extracellular matrix and seam cell shape (Figure 6). Failure of these tissues to properly compact and thicken due to a loss of DEX-1 could lead to an overall weakening of the cuticle and thus result in SDS sensitivity observed in *dex-1* mutant dauers. Interestingly, DEX-1 is similar to the human extracellular matrix protein SNED1 (Sushi, nidogen, and EGF-like Domain 1) that promotes invasiveness during breast cancer metastasis (Naba *et al.* 2014), suggestive of a mechanical role in tissue remodeling. Previous research demonstrated a role for autophagy dauer-specific seam cell remodeling (Melendez *et al.* 2003). It will be interesting to determine if dauer-specific changes to autophagy are influenced through DEX-1 mediated mechanical forces.

**Figure 6.**
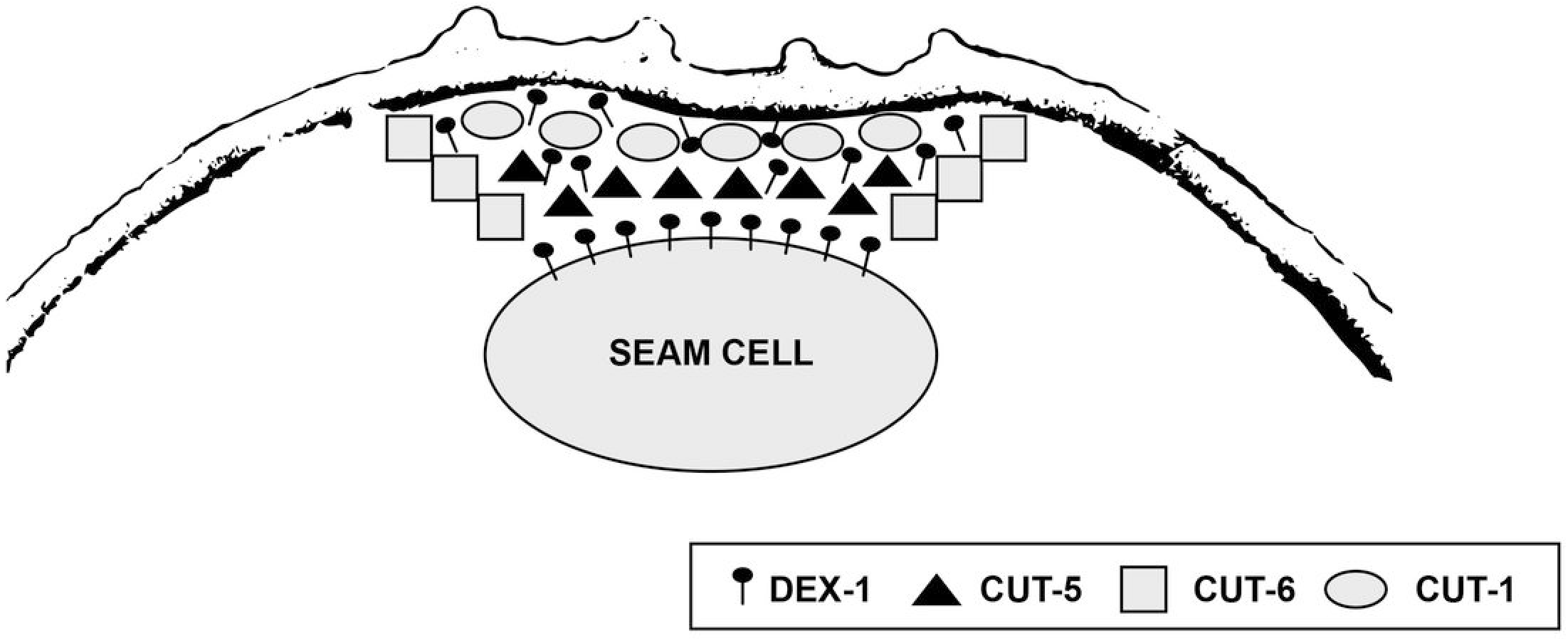
Models of DEX-1 interaction with cuticlin proteins. DEX-1 acts in a cell-autonomous manner along with the cuticlin proteins to facilitate alae formation. CUT-1 and CUT-5 are localized to ribbons directly below the alae, while CUT-6 localizes to the edges of the lateral cuticle (Muriel et al., 2003, Sapio et al., 2005). We propose that DEX-1 acts as a molecular anchor in the seam cell membrane to interact with multimerized cuticlin proteins in the cuticle. Interactions between DEX-1 and the cuticlins leads to compaction of the seam cell. Compaction of the seam leads to the formation of dauer-specific alae. Alternatively, DEX-1 may be secreted, but highly localized to the apical membrane of the seam cell. In this model, DEX-1 functions as a binding hub by facilitating interactions between the cuticlin proteins in the extracellular space.

## Acknowledgments

We thank Max Heiman for reagents and advice, and Scott Robinson at the Beckman Institute for assistance with electron microscopy. This work was supported by the National Institutes of Health (grant R01GM111566 to NES).

